# Multivariate association between single-nucleotide polymorphisms in Alzgene linkage regions and structural changes in the brain: discovery, refinement and validation

**DOI:** 10.1101/088310

**Authors:** Elena Szefer, Donghuan Lu, Farouk Nathoo, Mirza Faisal Beg, Jinko Graham, for the Alzheimers Disease Neuroimaging Initiative

**Affiliations:** Department of Statistics and Actuarial Science, Simon Fraser University, 8888 University Dr, Burnaby, BC, Canada V5A 1S6

## Abstract

Both genetic variants and brain region abnormalities are recognized to play a role in cognitive decline. We explore the association between singlenucleotide polymorphisms (SNPs) in linkage regions for Alzheimer’s disease and rates of decline in brain structure using data from the Alzheimers Disease Neuroimaging Initiative (ADNI).

In an initial discovery stage, we assessed the presence of linear association between the minor allele counts of 75,845 SNPs in the Alzgene linkage regions and predicted rates of change in structural MRI measurements for 56 brain regions using an RV test. In a second, refinement stage, we reduced the number of SNPs using a bootstrap-enhanced sparse canonical correlation analysis (SCCA) with a fixed tuning parameter. Each SNP was assigned an importance measure proportional to the number of times it was estimated to have a nonzero coefficient in repeated re-sampling from the ADNI-1 sample. We created refined lists of SNPs based on importance probabilities greater than 50% and 90%, respectively. In a third, validation stage, we assessed the multivariate association between these refined lists of SNPs and the rates of structural change in the independent ADNI-2 study dataset.

There was strong statistical evidence for linear association between the SNPs in the Alzgene linkage regions and the 56 imaging phenotypes in both the ADNI-1 and ADNI-2 samples (*p* < 0.0001). The bootstrap-enhanced SCCA identified 1,694 priority SNPs with importance probabilities > 50% and 22 SNPs with importance probabilities > 90%. The 1,694 prioritized SNPs in the ADNI-1 data were associated with imaging phenotypes in the ADNI-2 data (*p* = 0.0021).

This manuscript presents an analysis that addresses challenges in current imaging genetics studies such as biased sampling designs and highdimensional data with low-signal. Genes corresponding to priority SNPs having the highest contribution in the validation data have previously been implicated or hypothesized to be implicated in AD, including GCLC, IDE, and STAMBP1andFAS. We hypothesize that the effect sizes of the 1,694 SNPs in the priority set are likely small, but further investigation within this set may advance understanding of the missing heritability in late-onset Alzheimers disease. Multivariate analysis; Linkage regions; Imaging genetics; Endophenotypes; Inverse probability weighting; Variable importance probabilities

## 1 Introduction

Alzheimer’s disease (AD) is a neurodegenerative disorder causing cognitive impairment and memory loss. The estimated heritability of late-onset AD is 60%80% (Gatz et al., 2006), and the largest susceptibility allele is the ε4 allele of *APOE* (Corder et al., 1993), which may play a role in 20% to 25% of AD cases. Numerous studies have identified susceptibility genes which account for some of the missing heritability of AD, with many associated variants having been identified through genome-wide association studies (GWAS) (e.g. Beecham et al., 2009; Kamboh et al., 2012; Bertram et al., 2008). Apart from *APOE*, the associated variants have mostly had moderate or small effect sizes, suggesting that the remaining heritability of AD may be explained by many additional genetic variants of small effect.

Identifying susceptibility variants with small effect sizes in GWAS is challenging since strict multiple testing corrections are required to maintain a reasonable family-wise error rate. This analysis focuses on leveraging information from prior family of studies of AD (Hamshere et al., 2007; Butler et al., 2009), by looking for association in previously identified linkage regions reported on the Alzgene website (Biomedical Research Forum, 2013). Linkage regions for AD are genomic regions that tend to be co-inherited with AD in families. By definition, linkage regions include susceptibility genes that are co-transmitted with the disease. The regions currently identified from family studies of AD are large, however, since families contain relatively few transmissions. Further transmissions over multiple generations would provide more fine-grain information about the location of susceptibility genes. Previous studies have fine-mapped a single linkage region through association of AD with genetic variants in densely genotyped or sequenced regions (Fallin et al., 2010; Ertekin-Taner, 2003; Scott et al., 2000; Züchner et al., 2008), or have confirmed linkage to AD in genomic regions identified from GWAS (Anna et al., 2011). In this report, we aim to fine-map multiple linkage regions for AD through multivariate association of their SNPs to the rates of atrophy in brain regions affected by AD.

We analyze data from two phases of the Alzheimer’s Disease Neuroimaging Initiative (Mueller et al., 2005), ADNI-1 and ADNI-2, which are case-control studies of AD and mild-cognitive impairment. The rates of atrophy in brain regions affected by AD are so-called endophenotypes: observable traits that reflect disease progression. By investigating the joint association between the genomic variants and the neuroimaging endophenotypes, we use the information about disease progression to supervise the selection of single-nucleotide polymorphisms (SNPs). This multivariate approach to analysis stands in contrast to the commonly-used mass-univariate approach (for a review, see Nathoo et al., 2017), in which separate regressions are fit for each SNP, and the disease outcome is predicted by the minor allele counts. Simultaneous analysis of association is preferred because the reduced residual variation leads to (i) a clearer assessment of the signal from each SNP, (ii) increased power to detect signal, and (iii) a decreased false-positive rate (Hoggart et al., 2008). We also employ inverse probability weighting to account for the biased sampling design of the ADNI-1 and ADNI-2 studies, an aspect of analysis that has not been accounted for in many previous imaging genetics studies (Zhu et al., 2016).

Methods that explicitly account for gene structure have been proposed for analyzing the association between multiple imaging phenotypes and SNPs in candidate genes (e.g. Wang et al., 2011; Greenlaw et al., 2017). However, these methods become computationally intractable when analyzing data with tens of thousands of genotyped variants. To select SNPs associated with disease progression, we instead use sparse canonical correlation analysis (SCCA) to find a sparse linear combination of SNPs having maximal correlation with the imaging endophenotypes. Multiple penalty schemes have been proposed to implement the sparse estimation in SCCA (Parkhomenko et al., 2009; Witten et al., 2009; Lykou and Whittaker, 2010). We employ an SCCA implementation that estimates the sparse linear combinations by computing sparse approximations to the left singular vectors of the cross-correlation matrix of the SNP data and the neuroimaging endophenotype data (Parkhomenko et al., 2009). Sparsity is introduced through soft-thresholding of the coefficient estimates (Donoho and Johnstone, 1994), which has been noted (Chalise and Fridley, 2012) to be similar in implementation to a limiting form of the elastic-net (Zou and Hastie, 2005). A drawback of *𝓁*_1_-type penalties is that not all SNPs from an LD block of highly-correlated SNPs that are associated with the outcome will be selected into the model (Zou and Hastie, 2005). We prefer an elastic-net-like penalty over alternative implementations with *𝓁*_1_ penalties because it allows selection of all potentially associated SNPs regardless of the linkage-disequilibrium (LD) structure in the data.

We may think of SNP genotypes as a matrix *X* and imaging phenotypes as a matrix *Y* measured on the same *n* subjects. Robert and Escoufier (1976) showed that estimating the maximum correlation between linear combinations of *X* and *Y* in canonical correlation analysis is equivalent to estimating the linear combinations having the maximum RV coefficient, a measure of linear association between the multivariate datasets (Escoufier, 1973). As the squared correlation coefficient between the first canonical variates, the RV coefficient is well-suited for testing linear association in our context. We use a permutation test based on the RV coefficient to assess the association between the initial list of SNPs in *X* and the phenotypes in *Y*. A permutation test with the RV coefficient is preferred over a parametric hypothesis test since the permutation null distribution is computed under the same conditions as the observed RV coefficient, resulting in a valid hypothesis test. The outcome of this test is used to determine whether or not to proceed with a second refinement stage that reduces the number of SNPs by applying SCCA.

Tuning parameter selection can be difficult when data has low signal (Nathoo et al., 2016) and selection of the soft-thresholding parameter in SCCA is challenging in our context. Since the number of SNPs exceeds the sample size and many of the SNPs are expected to be unassociated with the phenotypes, large sample correlations can arise by chance (Fan et al., 2011). Indeed, the prescribed procedure of selecting the penalty parameter with highest predicted correlation across cross-validation test sets (Parkhomenko et al., 2009) results in more than 98% of the SNPs remaining in the model. A prediction criterion for choosing the penalty term may contribute to the lack of variable selection, allowing redundant variables into the model (Leng et al., 2006). When the same tuning parameter is used for variable selection and shrinkage, redundant variables tend to be selected to compensate for overshrinkage of coefficient estimates and losses in predictive ability (Radchenko and James, 2008). In our case, there is effectively no variable selection and little insight is gained by allowing for sparsity in the solution. To circumvent the lack of variable selection from SCCA, we fix the tuning parameter to select about 10% of the SNPs (Wu et al., 2009) and then use resampling to determine the relative importance of each SNP to the association with neuroimaging endophenotypes. Instead of using the prediction-optimal penalty term, we fixed the soft-thresholding parameter for the SNPs to achieve variable selection based on the rationale that no more than about 7,500 SNPs, or approximately 10%, are expected to be associated with the phenotypes. This choice is guided by prior experience in genetic association studies, where the majority of genetic variants have no effect on the phenotypes, or an effect that is indistinguishable from zero (Carbonetto and Stephens, 2012).

The organization of the manuscript is as follows. The Materials and Methods section describes the ADNI data, the data processing procedures, and the methods applied for discovery, refinement, and validation. The Results section presents the results of the analyses. The Discussion section notes challenges and successes of the analysis, including considerations for modelling continuous phenotype data under a case-control sampling design, and provides interpretation of the results.

## 2 Materials and Methods

### 2.1 Materials

Figure 1 illustrates the data processing steps required to compute the quantities that are analyzed from the raw data: adjusted minor allele counts of SNPs in the Alzgene linkage regions for the genomic data, and adjusted predicted rates of change at 56 brain regions of interest (ROIs) from MRIs for the neuroimaging data. The following subsections detail the data processing steps required to obtain the analysis and validation datasets.

**Figure 1:**
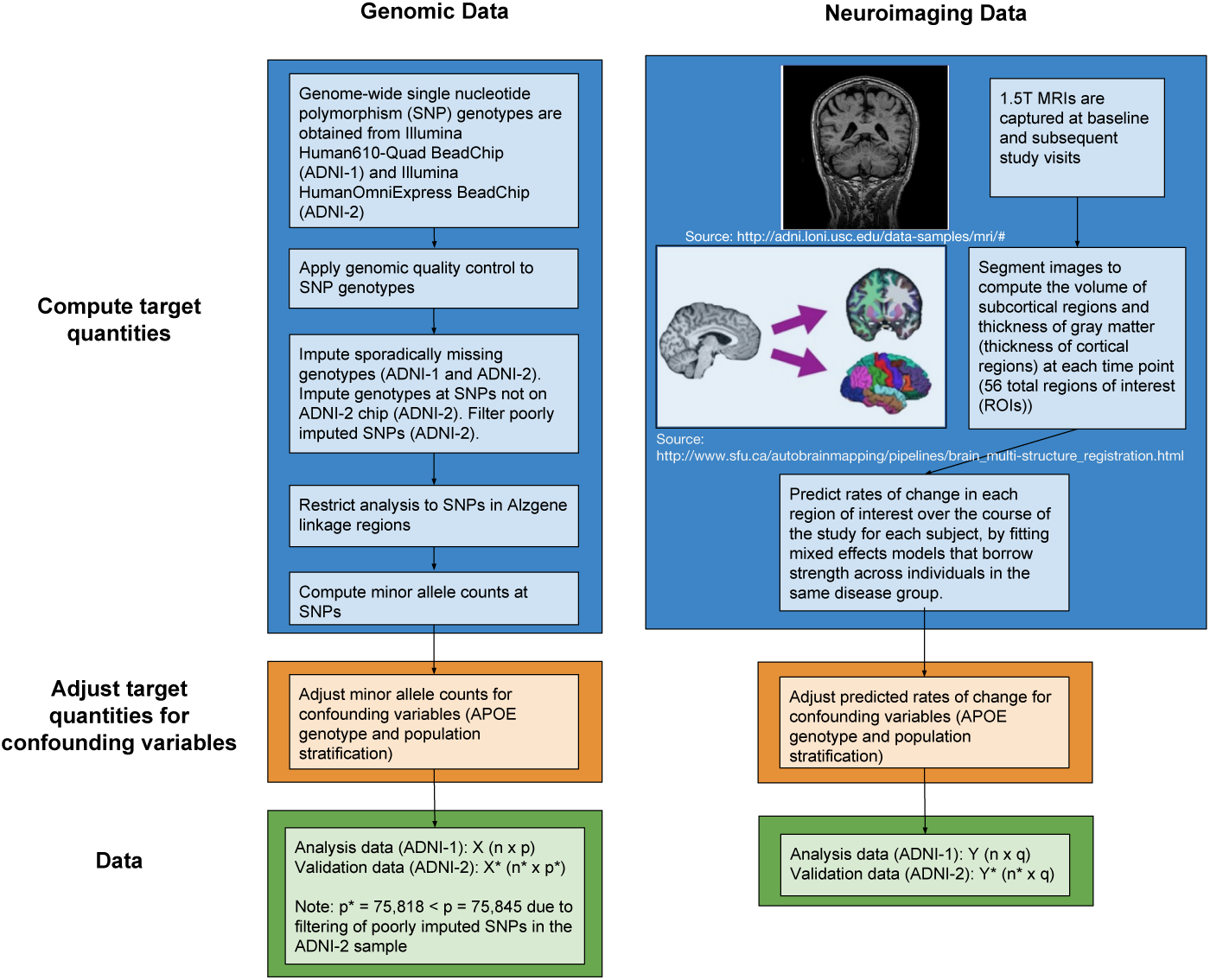
Flow chart of the steps to process the raw genomic and neuroimaging data into the analysis and validation datasets. The target quantities, minor allele counts at SNPs in Alzgene linkage regions for the genomic data and predicted rates of change at brain regions of interest for the neuroimaging data, are computed. Then, both sets of targets quantities are adjusted for potentially confounding variables to obtain the data for analysis (ADNI-1) or validation (ADNI-2).

#### 2.1.1 ADNI

Data used in the preparation of this article were obtained from the Alzheimer’s Disease Neuroimaging Initiative (ADNI) database (adni.loni.usc.edu). The primary goal of ADNI has been to test whether serial magnetic resonance imaging (MRI), positron emission tomography (PET), other biological markers, and clinical and neuropsychological assessment can be combined to measure the progression of mild cognitive impairment (MCI) and early Alzheimer’s disease (AD). Patients with MCI have a subjective memory concern but no significant cognitive impairment, while patients with early Alzheimer’s disease have experienced memory loss. The first phase of ADNI, ADNI-1, aimed to enroll approximately 800 study subjects, aged 55-90, of which 200 were cognitively normal (CN), 200 had mild Alzheimers disease (AD), and 400 had mild cognitive impairment (MCI) (ADNI Procedures Manual, 2006, p. 3). The CN subjects were roughly age-matched to the MCI and AD subjects. Subjects were assigned to a group following inclusion and exclusion criteria based on clinical and cognitive tests. Mild AD subjects had mini-mental state examination (MMSE) scores from 20-26 inclusive, clinical dementia rating (CDR) of 0.5 or 1.0, and met the National Institute of Neurological and Communicative Disorders and Stroke and the Alzheimers Disease and Related Disorders Association (NINCDS/ADRDA) criteria for probable AD. MCI subjects had MMSE scores between 24-30, CDR of 0.5, a memory complaint, and objective memory loss measured by education adjusted scores on Wechsler Memory Scale Logical Memory II (Wechsler, 2009). Cognitively normal participants were non-depressed, with MMSE scores between 24-30, and did not fit the criteria for the MCI or mild AD groups (ADNI Procedures Manual, 2006, p. 3-4). The study subjects were enrolled from 50 sites in the US and Canada and were required to commit to at least two years of follow up MRI.

ADNI-2 was a subsequent phase of ADNI that involved enrollment of new subject cohorts, as well as continued follow-up of subjects enrolled in earlier phases. ADNI-2 enrolled a new cohort of subjects among which 190 subjects were cognitively normal, 181 subjects had early MCI, 164 subjects had MCI, and 148 subjects had mild AD.

The ADNI-1 cohort was used in our initial analysis. To validate the findings of the initial analysis, a subset of the ADNI-2 cohort was used that included new subjects in the same disease categories (CN, MCI and AD) as the subjects in ADNI-1.

#### 2.1.2 Imaging data

The neuroimaging phenotypes analyzed are predicted rates of change in cortical thickness and volumetric measurements in brain regions obtained from magnetic resonance imaging (MRI) scans. ADNI subjects had 1.5T MRI scans at either 6 or 12 month intervals during the two- to three-year follow-up period of the study and we chose to analyse the longitudinal information on cortical thickness and regional volumes. While other studies have compared the different study groups using imaging information from baseline (Shen et al., 2010; Meda et al., 2012), the longitudinal information provides insight into the different rates of brain deterioration experienced by people with negligible memory loss compared to those with more acute memory difficulties and Alzheimer’s disease.

##### Segmentation

Segmentation is the process of identifying the locations of anatomical structures within an image. In these data, the MRIs were segmented using Freesurfer (Fischl, 2012) software, identifying the locations of regions of interest such as the hippocampus, cerebellum and ventricles. For each hemisphere, the 28 volumetric and cortical thickness measurements used for analysis by Shen et al. (2010) were obtained via automated parcellation of the segmented images in Freesurfer. Cortical thickness (thickness of gray matter) and volumes of the regions of interest become increasingly atrophied as disease progresses, so we expect increased rates of atrophy in participants with more memory concerns (the MCI and AD subjects) compared to the cognitively normal subjects.

##### Predicting rates of change

Linear mixed effect models, given in Equation 1, were used to predict the rates of change in each brain region of interest (ROI). Brief descriptions of the regions of interest are given in Appendix A. A separate mixed model was fit for each ROI, with random effects for subject-specific rates of change and fixed effects for average rates of change within diagnostic subgroups. The response variable *Y*_*i* *jt*_ is a continuous measurement of the cortical thickness or volume of a brain region of interest. In the specification of the model, fixed-effects terms are denoted by *β*, while random-effect terms are denoted by γ. The covariates are (i) *t*, the time of the follow-up visit at which the scan was conducted, *∈* with *t* ∈ 0, 6, 12, 18, 24 months; (ii) *MCI*, a dummy variable equal to 1 if subject *i* has mild cognitive impairment, and equal to 0 otherwise; and (iii) *AD*, a dummy variable equal to 1 if subject *i* has Alzheimer’s disease, and equal to 1 otherwise. The ROI’s are indexed by *j*:

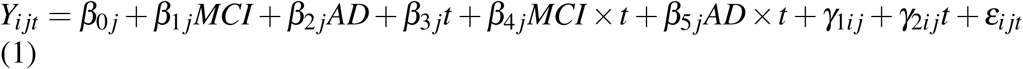

The predicted rate of change over the study period for subject *i* at ROI *j* is the sum of the disease-specific estimated rate of change and the subject-specific predicted rate of change 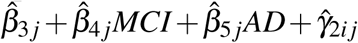 The lme4 package (Bates et al., 2015) was used to fit the mixed effects model in R, and the conditional mode was used to predict the random effect γ_2*i* *j*_. The normality of the predicted, subject-specific, random effects was assessed by examining the Q-Q plots of the predictions at each ROI. The assumption of normality appeared to be reasonable for all ROIs.

Figure 2 is a heatmap of the predicted rates of change, adjusted for potential confounding variables in the sample, as discussed next. The heatmap illustrates how rates of change are more negative in subjects with more advanced disease, indicating that the thickness of gray matter and the volumes of brain regions of interest are shrinking more. Decreases in cortical thickness are more pronounced for subjects with AD (i.e. their rates of changes are more negative for various cortical thickness measures). Similarly, the ventricles, cavities in the brain filled with cerebrospinal fluid, have a more positive rate of change for subjects with more advanced disease. As brain atrophy progresses, these cavities expand.

#### 2.1.3 Genomic data

The ADNI-1 subjects were genotyped with the Illumina Human610-Quad BeadChip and the ADNI-2 subjects were genotyped with the Illumina HumanOmniExpress BeadChip, both of which interrogate SNPs. All genotyping information was downloaded from the LONI Image Data Archive (Laboratory of Neuroimaging, 2015). Pre-packaged PLINK (Purcell et al., 2007) files included genotyping information for 757 of the 818 ADNI-1 subjects. Genotyping information for 793 of the ADNI-2 subjects were converted from CSV files to PLINK binary files using a publicly-available conversion script (Hibar, 2014). The Human610-Quad BeadChip and HumanOmniExpress BeadChip interrogated 620,901 and 730,525 SNPs respectively. APOE was genotyped separately at study screening from DNA extracted from a blood sample.

##### Inclusion criteria

Subjects were included if their genotyping data were available, if they had a baseline MRI scan, and they had at least one additional follow-up baseline scan. Of the 757 ADNI-1 subjects and 793 ADNI-2 subjects with SNP data available, 696 and 583 had both a baseline scan and at least one additional follow-up scan, respectively. Genomic quality control procedures outlined below in the Genomic quality control section were also applied, widening the exclusion criteria, to obtain a more homogeneous sample.

**Figure 2:**
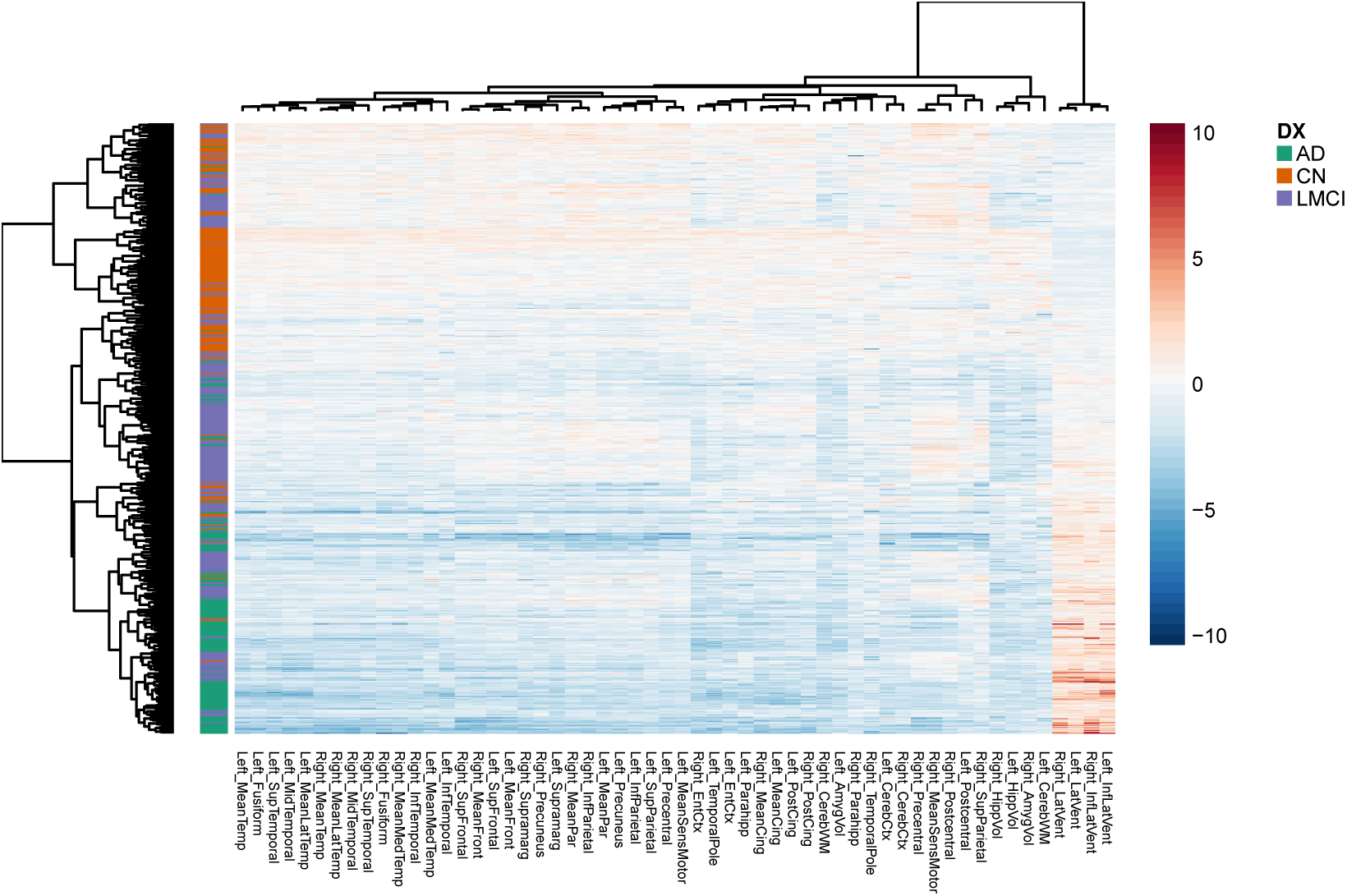
Heatmap of the neuroimaging phenotypes, clustered by similarity among ROIs and subjects. Each row corresponds to a subject in the sample and each column corresponds to one of the 56 ROIs. The rows are annotated by the disease group of the subject. The adjusted, predicted rates of change are shown for each region, where blue values indicate decreases in the volume of thickness in the brain region, and orange values indicate increases in the volume of the brain region. Values for the ventricles clustered on the far right, have an inverted relationships compared to the other ROIs since the ventricles are cavities in the brain which expand as brain atrophy progresses. The thickness of gray matter, by contrast, decreases with atrophy.

##### Genomic quality control

Genomic quality control was performed with the freely-available software package PLINK. Thresholds for SNP and subject exclusion were as defined in Shen et al. (2010), except that we required a more conservative genotyping call rate for SNPs, as in Ge et al. (2012). Quality control was applied in three separate phases:

- Phase 1

1. Exclude SNPs with genotyping call rate< 95%
2. Restrict sample to subjects with self-reported non-Hispanic Caucasian ethnicity and white race.

- Phase 2

1. Exclude SNPs with minor allele frequency (MAF) < 5%, Hardy-Weinberg equilibrium (HWE) *p* < 10^−6^.
2. Exclude subjects based on tests for multivariate outliers and tests of relationship and gender using the genotyping data.

- Phase 3

1. Exclude SNPs from sex chromosomes.

Details of the genomic quality control procedures that were applied may be found in Szefer (2014).

##### Genomic imputation

Imputation serves two key roles in the analysis: to preserve the sample size for the multivariate analysis by replacing sporadically missing genotypes with imputed ones, and to impute SNPs not interrogated on the ADNI-2 chip that are interrogated on the ADNI-1 chip. Best-guess SNP genotypes were imputed in the ADNI-1 and ADNI-2 sample using the HapMap3 panel with NCBI build 36/hg18 using IMPUTE2 (Marchini and Howie, 2010), based on the imputation protocol in the IMPUTE2: 1000 Genomes Imputation Cookbook (Luan et al., 2014). Haplotypes were phased with SHAPEIT (Delaneau et al., 2013), and PLINK and SHAPEIT/IMPUTE2 file formats were converted with GTOOL (Freeman, 2007–2012). Out of the 503,450 SNPs that passed quality control in the ADNI-1 sample, sporadically missing genotypes were imputed at the 459,517 SNPs that were also in the reference panel. Out of the 574,730 SNPs that passed quality control in the ADNI-2 sample, sporadically missing genotypes were imputed at the 270,074 SNPs that were also on the ADNI-1 chip and in the reference panel. The remaining 189,443 SNPs that were not genotyped in the ADNI-2 sample, but were in both the ADNI-1 sample and the reference panel, were imputed into the sample. The genotyping rate in the imputed data for the ADNI-2 sample was 98.2%, prior to filtering out SNPs with an IMPUTE2 info metric < 0.5. The IMPUTE2 metric measures the reliability of imputed genotypes for a SNP, and takes a value of 1 when there is no genotype uncertainty in the sample. When the metric is less than 0.5, the sample mean of the posterior variance of imputed genotypes is at least half the variance that would be expected if alleles were sampled at random (Marchini and Howie, 2010).

The quality of the imputation for a SNP is reported based on the IMPUTE2 certainty measure, which is the average posterior probability of the best-guess genotypes in the sample. In the ADNI-1 data, the average certainty was 100% for all SNPs in the Alzgene linkage regions. In the ADNI-2 data, the certainties ranged from 69.1% to 100%, but 98.4% of the 31,301 imputed SNPs in the Alzgene linkage regions (defined below) had certainties of 90% or more. Due to the scale of the analysis, the imputed genotypes were treated as known. Ignoring genotype uncertainty is expected to lead to underestimates of the variance in the analysis of the ADNI-2 but not the ADNI-1 data.

##### Alzgene linkage regions

To focus the analysis on regions that are likely to contain causal genetic variation, SNPs were included in the analysis if they fell in the linkage regions reported by approximate physical position on the Alzgene website (Bertram et al., 2007; Biomedical Research Forum, 2013). These linkage regions have been identified in meta-analyses of family-based studies of Alzheimer’s disease (Hamshere et al., 2007; Butler et al., 2009). A total of 75,845 SNPs from nine chromosomes were included in the analysis from the ADNI-1 sample. Table 1 shows the number of SNPs in the ADNI-1 sample that fall in each linkage region. After filtering SNPs that had an IMPUTE2 info metric < 0.5, 75,818 SNPs remained in the ADNI-2 sample.

#### 2.1.4 Adjustments for confounding

Covariate information cannot be explicitly included in SCCA, so both the imaging and genomic data are adjusted for confounding variables in advance. Potential confounders in the analysis are population stratification and APOE genotype. Population stratification is the phenomenon of systematic differences in allele frequencies in a subpopulation arising because of differences in ancestry, while the ε4 allele of APOE is the largest known genetic risk factor for Alzheimer’s disease (Corder et al., 1993). Since true population structure is not observed, we adjust for it in the data using the top ten principal coordinates from multidimensional scaling. The top 10 principal coordinates account for 2.6% of the variability in the genotype data. We also restrict the analysis to the white non-Hispanic subjects and remove multivariate outliers identified in the top two principal coordinates as noted in the Genomic quality control section. We adjust for APOE genotype as a precautionary measure, since it can account for the population stratification in the data, over and above the principal components or principal coordinates (Lucotte et al., 1997).

**Table 1:** The chromosome, band, and location on the Mb scale of the linkage regions of interest. *N* denotes the number of SNPs in the ADNI-1 data that fall in each linkage region.

| Chromosome | Band | Mb | *N* |
| --- | --- | --- | --- |
| 1 | p31.1-q31.1 | 83-185 | 12005 |
| 3 | q12.3-q25.31 | 103-173 | 10689 |
| 6 | p21.1-q15 | 43-91 | 6785 |
| 7 | pter-q21.11 | 0-78 | 13292 |
| 8 | p22-p21.1 | 13-28 | 4149 |
| 9 | p22.3-p13.3 | 20-35 | 2868 |
| 9 | q21.31-q32 | 80-100 | 3483 |
| 10 | p14-q24 | 10-100 | 15274 |
| 17 | q24.3-qter | 67-79 | 2319 |
| 19 | p13.3-qter | 8-54 | 4981 |

Ten principal coordinates for each of the ADNI-1 and ADNI-2 datasets were obtained using ten-dimensional multi-dimensional scaling on the pairwise IBS distance matrix, computed with PLINK from 121,795 and 118,012 approximately uncorrelated SNPs from the SNPs that passed quality control filters. The SNP genotypes used to estimate the principal coordinates were from the complete imputed data. The number of principal coordinate dimensions was chosen to follow a similar protocol for adjustment for population stratification using principal components, in which ten axes of variation are suggested (Price et al., 2006).

The data for analysis were obtained by adjusting the minor allele counts and predicted rates of change of the brain ROIs for the ten principal coordinates, as well as for dummy variables for APOE genotype, using weighted ordinary least squares regression. The weights account for certain diagnostic subgroups being over-represented in the sample relative to their population frequency. Details on the computation of the weights are presented in the next section. The residuals from each regression comprised the genomic (*X*) and neuroimaging (*Y*) features analyzed.

### 2.2 Methods

Figure 3 illustrates each step in the data analysis process, from deriving inverse probability weights to account for the biased sampling in ADNI-1 and ADNI2, to discovering, refining and validating association between the adjusted minor allele counts at SNPs, which we call the genotypes, and the adjusted predicted rates of change at the brain regions of interest, which we call the neuroimaging phenotypes.

#### 2.2.1 Inverse probability weights

To account for the biased sampling in the ADNI-1 and ADNI-2 case-control studies, we estimated inverse probability weights for each subject (Horvitz and Thompson, 1952). As subjects with early MCI were excluded from ADNI-1, we defined the target population to be non-Hispanic, white Americans and Canadians aged 55-90 years who are cognitively normal or have been diagnosed with late MCI or Alzheimer’s disease.

The Alzheimer’s Association reports that 5.2 million Americans had Alzheimer’s disease in 2014 (Alzheimer’s Association, 2014). Additionally, data from the US census in 2010 (U.S. Census Bureau, 2011) indicate that approximately 23% of the American population is over the age of 55 and that the total population is 308 million people. Based on this information, the approximate proportion of the American population aged 55-90 years with Alzheimer’s disease is *p*_*AD*_ = 7.5%, rounded to the nearest half percent. This calculation assumes that individuals aged 90 or more years and patients diagnosed with early MCI represent negligible proportions of the population. We used a late MCI prevalence estimate of *p*_*MCI*_ = 5% based on an urban study of people aged 65+ in New York (Manly et al., 2005), and assumed that the remaining *p*_*CN*_ = 87.5% of the population of interest is cognitively normal. A breakdown of the number of subjects used in the analysis by study is given in Table 2.

The inverse probability weights for each disease group, *w*_*DX*_ for ADNI-1 and 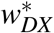 for ADNI-2, are computed as the assumed prevalence of the disease in the target population divided by the number of subjects sampled from the disease group. In the ADNI-1 sample, the MCI subjects have *w*_*MCI*_ = 0.11, AD subjects have *w*_*AD*_ = 0.30 and CN subjects have *w*_*CN*_ = 3.09, where the weights have been standardized to sum to the ADNI-1 sample size of *n* = 632. In the ADNI-2 sample, the MCI subjects have 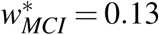, the AD subjects have 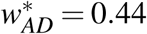 and the CN subjects have 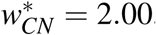, where the weights have been standardized to sum to the ADNI-2 sample size of *n* = 265.

**Figure 3:**
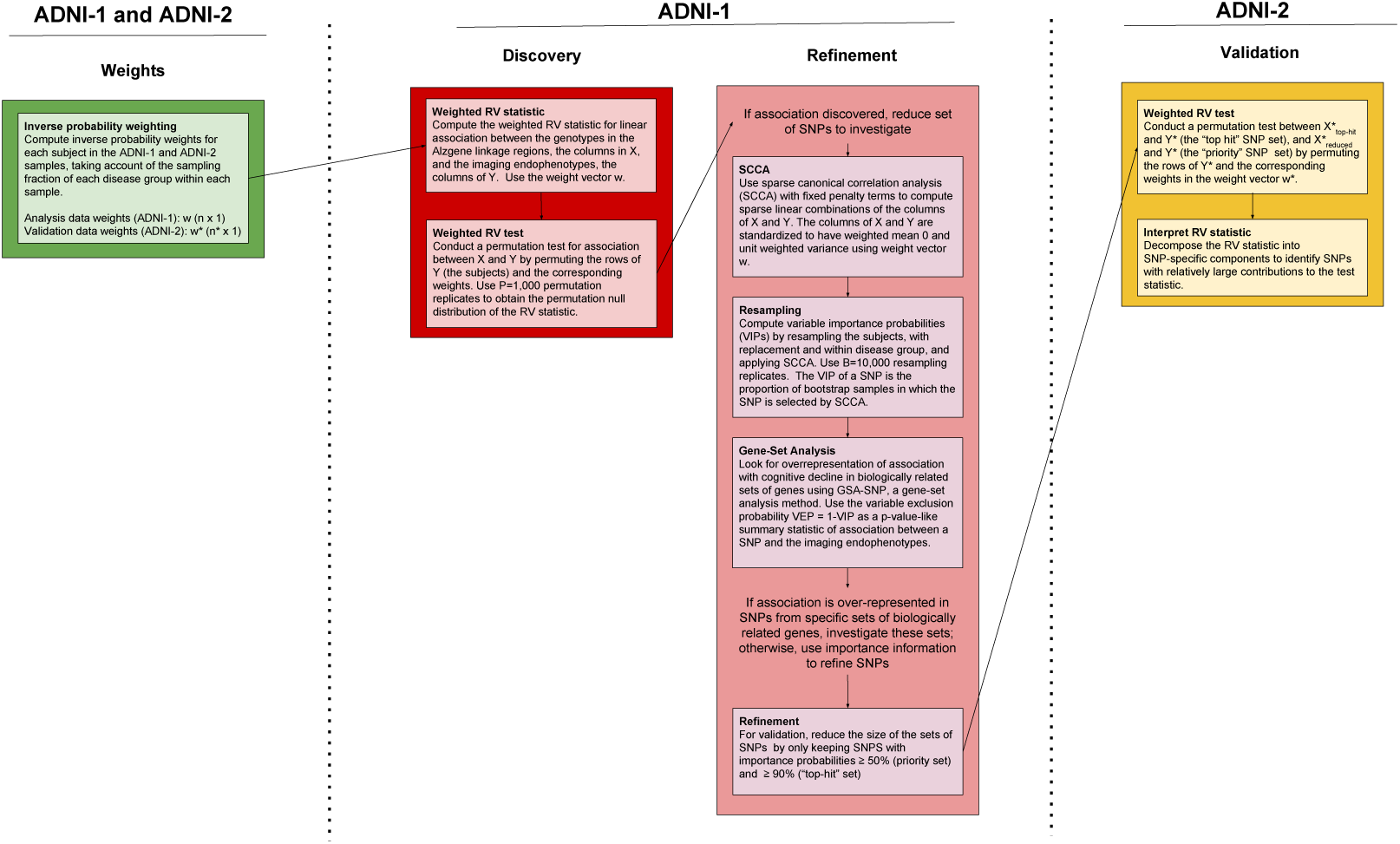
Flow chart of the analysis steps. The first row of column headings indicate the data sample used in the analysis step. The second row of column headings denote the step in the analysis beginning with computing weights, discovering association, refining the set of SNPs to investigate, and ending with validation of association with refined sets of SNPs and the neuroimaging phenotypes.

**Table 2:** The number of subjects, *n*_*D*_, from each disease group *D* that were analyzed in each study. The total number of subjects analyzed in each study is denoted by *n*.

| Sample | $n_{CN}$ | $n_{MCI}$ | $n_{AD}$ | $n$ |
| --- | --- | --- | --- | --- |
| ADNI-1 | 179 | 296 | 157 | 632 |
| ADNI-2 | 116 | 104 | 45 | 265 |

#### 2.2.2 Discovery

##### Weighted RV test

We tested the analysis dataset, ADNI-1, for linear association between the genomic data and the neuroimaging data. The RV coefficient (Escoufier, 1973) is a multivariate generalization of Pearson’s *r*^2^ and quantifies the association between the columns of *X*, or the genotypes, and the columns of *Y*, the imaging endophenotypes. The coefficient can be defined in terms of the sample covariance matrices *S*_*XX*_ and *S*_*YY*_, and the cross covariance matrix *S*_*XY*_(Omelka and Hudecova, 2013), where the (*k, l*)th element of *S*_*XY*_ is defined as:

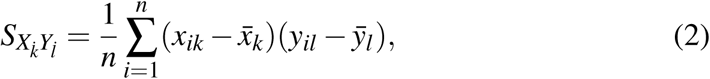

Where 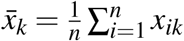 and 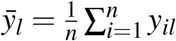 The (*k, l*)th elements of *S*_*XX*_ and *S*_*YY*_ are defined similarly. The RV coefficient is then written as:

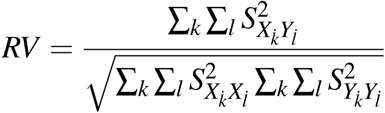

and takes values between 0 and 1. When *RV* = 0, there is no linear association between the columns of *X* and *Y*, and higher values of *RV* are indicative of more association.

As the test statistic, we used a weighted version of the RV coefficient (Omelka and Hudecova, 2013) that calculates the sample variances and cross covariances accounting for the oversampling of AD and MCI patients in the study. A permutation test with P=10,000 permutations was used to assess the evidence for association between *X* and *Y*, where the rows of *Y* and their associated inverse probability weights were randomly permuted.

#### 2.2.3 Refinement

##### SCCA and resampling

Sparse canonical correlation analysis (SCCA; Parkhomenko et al., 2009) is a multivariate method for estimating maximally correlated sparse linear combinations of the columns of two multivariate datasets collected on the same *n* subjects, *X* and *Y*. To obtain a sparse linear combination of the SNP genotypes that is most associated with a non-sparse linear combination of the imaging phenotypes, we used SCCA, a penalized version of canonical correlation analysis. Sparse linear combinations contain some coefficients which are zero; e.g., in penalized regression analysis, the predicted value is potentially a sparse linear combination of the predictors. SCCA operates on the cross-correlation matrix, which is equivalent to the cross-covariance matrix *S*_*XY*_ as defined in Equation 2 when the columns of *X* and *Y* are standardized. SCCA estimates sparse linear combinations *aX* and *bY* that have maximal correlation, where *a* and *b* are column vectors of length *p* and *q* respectively. By operating on the cross-correlation matrix, the distance metric used is Euclidean, which is appropriate since both the columns of *X* and *Y* are treated as continuous in the analysis; other distance metrics for the genetic data are possible, however (Chang, 2017).

We initially applied SCCA to identify a sparse set of SNPs associated with the imaging endophenotypes. Ten-fold cross validation was used to select the penalty parameter for the SNPs, λ_*u*_, for the SCCA. A search grid for λ_*u*_ was defined as {0, 10^−4^,…, 10^−1^} with the values in the search grid being incremented by 0.0005. At the *i*^*th*^ element in the search grid, λ_*u,i*_, the sparse canonical-correlation coefficients, *a*_*i,* *j*_ for the SNPs and *b*_*i,* *j*_ for the endophenotypes, were computed in training set *j*. The fitted coefficients from the training sets were then used to compute the predicted sample-correlation coefficient in each test set: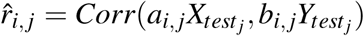. The SNP penalty parameter λ_*u*_ was chosen as the element in the search grid that maximized the sum of the predicted sample-correlation coefficients over the ten test sets. Under this cross-validation scheme, variable selection of the SNPs was minimal with more than 98% of the SNPs remaining in the active set. Ruling out fewer than 2% of the SNPs in the Alzgene linkage regions is insufficient refinement for our analysis.

Instead, we chose to incorporate bootstrap resampling to estimate the relative importance of each SNP in the multivariate association. This approach of “bootstrap enhancement” has been applied previously in neuroimaging studies (Bunea et al., 2011), to guide variable selection with the elastic-net and the lasso. We obtained B=100,000 bootstrap samples by sampling subjects with replacement within each disease category. The weighted cross-correlation matrix 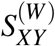 was computed for each bootstrap sample *b*, and a sparse linear combination of the genomic markers was estimated, using the SCCA penalty parameter 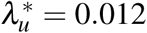 for soft-thresholding the SNP coefficients. A value of 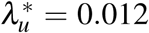 was chosen so that approximately 10% of the SNPs had non-zero estimated coefficients. If β _*b*_ = (β_1*b*_, β_2*b*_,…, β_*pb*_) denotes the coefficient vector of the sparse linear combination of the *p* SNPs, from bootstrap sample *b*, then the variable importance probability for SNP *k* (*VIP*_*k*_) is defined as the proportion of bootstrap samples in which SNP *k* (*k* = 1, *p*) has a nonzero coefficient, or is “selected”:

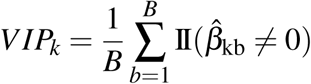, where II(A) = 1 if condition A holds and 0 otherwise

##### Gene-Set Analysis

To reduce the initial list of 75,845 SNPs to a shorter list for validation and to gain insight into the biologically related sets of genes associated with cognitive decline, we applied a gene-set analysis, as implemented in GSA-SNP (Nam et al., 2010). GSA-SNP combines the evidence for SNP-specific associations into gene-level summaries and assesses the pattern of association for genes in a given set, such as a functional pathway, relative to genes outside the set We used variable exclusion probabilities, *VEP* = 1*-VIP*, to quantify the SNP-specific evidence of association in lieu of p-values, and the second smallest *VEP* for SNPs in a gene as the gene-level summary statistic. Our choice of the second smallest was based on the recommendation of Nam et al. (2010) to use the second smallest rather than the smallest p-value as the gene-level summary statistic to protect against the spuriously high association signal that may be introduced by longer genes. The re-standardized version of GSA-SNP with the maxmean statistic (Efron and Tibshirani, 2007) was applied, with default gene padding of 20000 base pairs and Gene Ontology gene sets. We took *P* = 100 samples under the permutation null hypothesis of no association to serve as the empirical reference distribution for *VEP*s from ADNI-1. To ensure inclusive selection of SNPs, candidate gene sets were identified by Benjamini-Hochberg corrected *p*-values with a liberal threshold of 0.8 for the false discovery rate.

##### Refinement

Two subsets of the SNPs in the Alzgene linkage regions, with estimated importance probabilities ≥ 50% and 90%, were used for validation in the ADNI-2 sample. The cut-off values were chosen to reflect a relatively liberal and stringent criterion, respectively. The set of SNPs with *VIP* ≥ 50% is called the “priority set”, while the SNPs with *VIP* ≥ 90% is called the “top-hit” set.

#### 2.2.4 Validation

##### Validation

We assessed the evidence for linear association between the tophit and priority SNPs and all the imaging phenotypes in the ADNI-2 validation sample using the RV-test with 1,000 permutation replicates.

##### Interpretation of RV statistic

To further understand the observed association between SNPs in the priority set and endophenotypes in the ADNI-2 validation data, we decomposed the RV test statistic into its SNP-specific components. Details on how the SNP-specific contributions are calculated are described in the Results section.

## 3 Results

### 3.1 Discovery

The RV-test in the ADNI-1 data rejected the null hypothesis of no linear association between *X* and *Y*. The observed RV coefficient was *RV* = 0.079, and the permutation test *p*-value was *p* < 0.001.

### 3.2 Refinement

The resampling procedure coupled with SCCA in the ADNI-1 data produces variable importance probabilities (VIPs) for each SNP in the Alzgene linkage regions. Figure 4 is a Manhattan-like plot of the variable exclusion probabilities, *VEP* = 1 -*VIP*, plotted on the −log_10_scale, such that SNPs with *VIP* ≥ 0.9, have values of −log_10_(*VEP*) ≥ 1. The dashed and dotted reference lines indicate the *VIP* = 0.5 and *VIP* = 0.9 cut-offs used to identify the priority and top-hit sets of SNPs, respectively.

1, 694 SNPs had *VIP* ≥ 0.5, a set of reduced SNPs we call the priority set. As expected, the priority SNPs, *X*_*reduced*_, were associated with the endophenotypes in the ADNI-1 training data, based on a permutation *RV* test (*RV* = 0.23). Using the stringent cut-off of *VIP* ≥ 0.9 for SNP selection, 22 SNPs were included in the “top-hit” set. There was no evidence of enrichment in biological pathways based on results from GSA-SNP.

**Figure 4:**
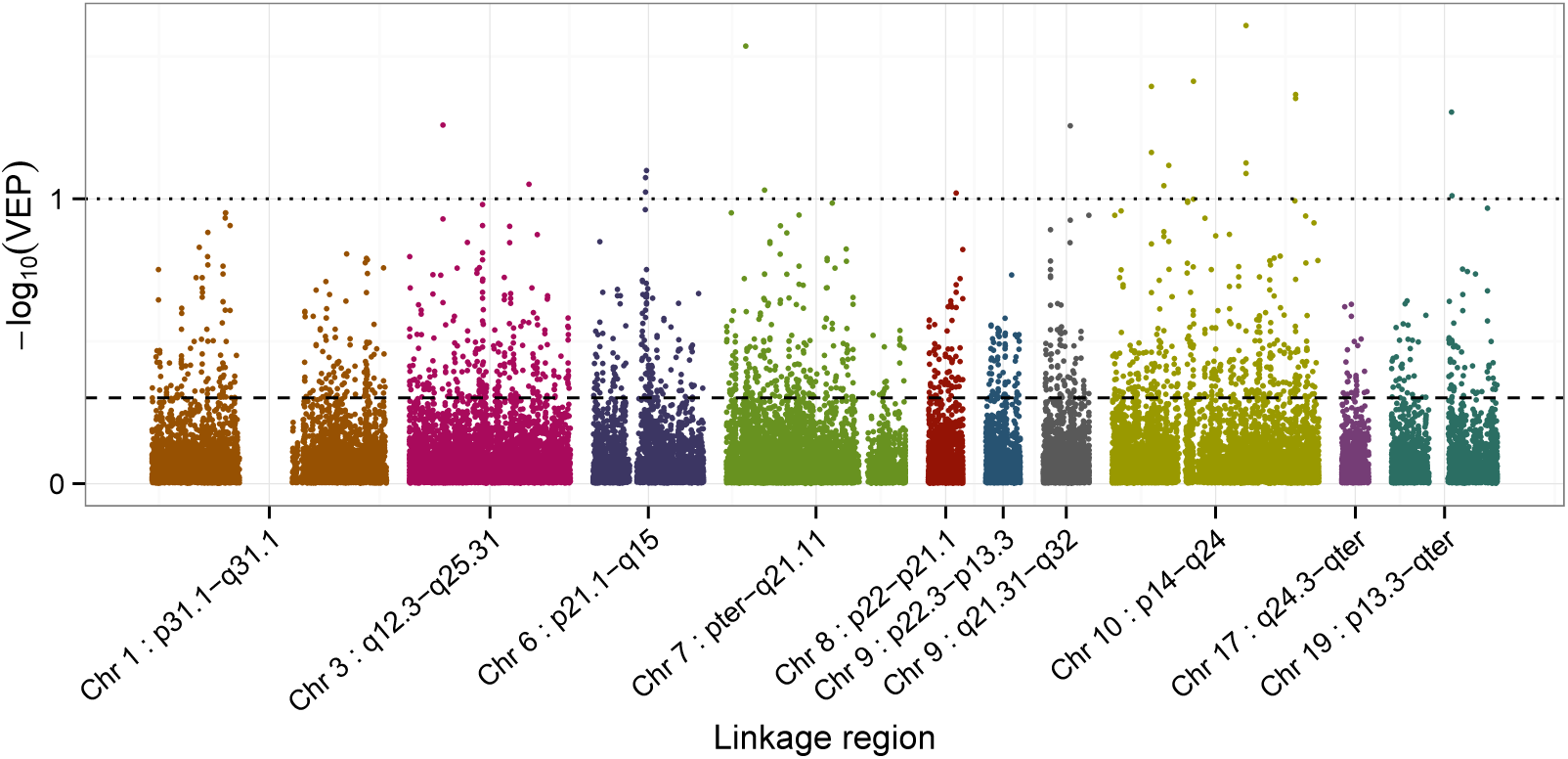
Plot of the –log_10_(*VEP*) of the SNPs in each of the Alzgene linkage regions. The dashed and dotted reference lines indicate the *VIP* = 0.5 and *VIP* = 0.9 cut-offs used to define the priority and top-hit sets of SNPs, respectively.

Figure 4 shows that very few SNPs had *VIP* ≥ 0.9, as evidenced by the sparse selection of SNPs with –log_10_(*VEP*) ≥ 1 in the plot. While the linkage region on chromosome 10 is the largest, it also has the most SNPs with *VIP* ≥ 0.9 and its SNPs have relatively high inclusion probabilities across the entire linkage region, in contrast to the linkage region from chromosome 6, for example. The smaller linkage regions p22.3-p-13.3 on chromosome 9 and q24.3-qter on chromosome 17 have relatively low inclusion probabilities, overall.

#### 3.3 Validation

Let 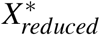 and 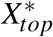 be the ADNI-2 validation data at the priority and top-hit sets of SNPs, respectively, and let *Y*\* be the validation endophenotype data. We were able to validate our finding of association between the priority set of SNPs and the endophenotypes in the ADNI-2 data. The *RV* test of association between 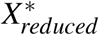 and *Y*\* had an observed test statistic of *RV*_*obs*_ = 0.073, and a permutation *p*-value of *p* = 0.0021. However, there was no evidence of association between the top-hit set of SNPs, 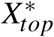, and the endophenotypes (*p* = 0.79).

Figure 5 depicts the contribution of each SNP as a score, normalized to have mean 1 over all the SNPs in the priority set, to the RV statistic. Before normalization, the contribution for SNP *i* in the priority set is a sum, 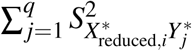 over the *q* = 56 endophenotypes in the cross-correlation matrix, 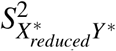 from the ADNI-2 validation data. Each point in the plot therefore represents the relative contribution of a given SNP to the *RV* coefficient, summed over the 56 endophenotypes. SNPs with higher relative contributions can be viewed as the SNPs driving the association found in the *RV* test.

**Figure 5:**
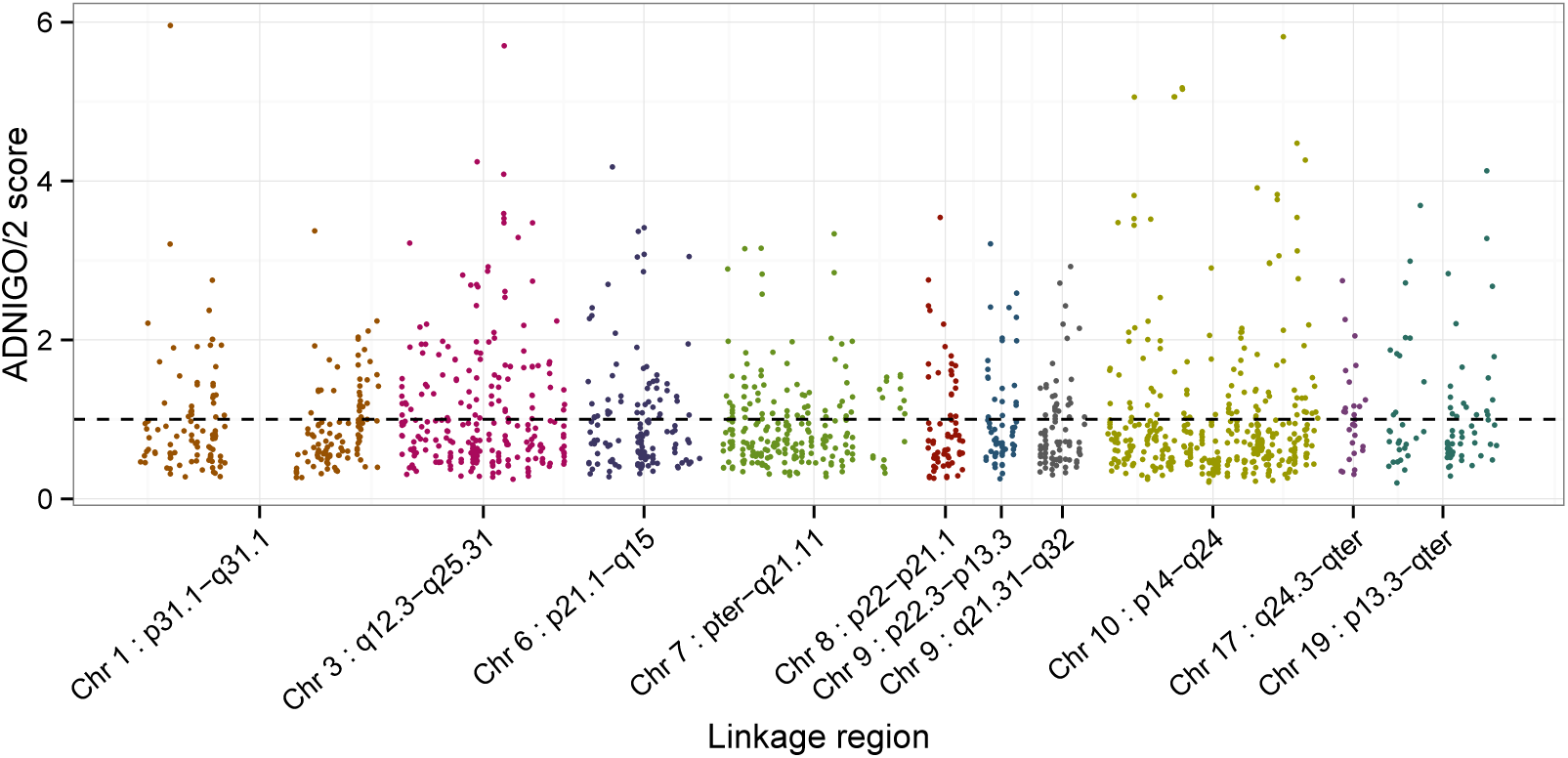
SNP-specific scores at the priority set SNPs in the ADNI-2 validation data, with scores defined as described in text. SNPs with higher score contribute relatively more to the *RV* coefficient between 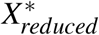 and *Y*\*. The dashed horizontal reference line corresponds to a score of 1, or the average score for a SNP in the priority set in the ADNI-2 validation data.

Table 3 summarizes information about the top 20 scoring SNPs in the priority set, with gene annotations obtained from SNPNexus (Ullah et al., 2012). SNPNexus was queried using assembly NCBI36/hg18, the UCSC genome browser (Speir et al., 2015) and AceView (Thierry-Mieg and Thierry-Mieg, 2006). The resulting gene symbols for annotated SNPs are reported in the Genes column of the table. We used the squared Pearson correlation coefficient, *r*^2^, to measure the linkage disequilibrium (LD) between SNPs. Values of *r*^2^ were computed in R with the snpMatrix package (Clayton and Leung, 2007) using the *N* = 116 cognitively normal subjects in the ADNI-2 data. LD blocks within the priority set are indicated by numbers in the first column of Table 3, and are defined such that all SNPs within a block have pairwise *r*^2^ greater than 0.7.

**Table 3:** The 20 SNPs with highest SNPs scores in the ADNI-2 dataset. Gene annotation obtained from SNPNexus queried with the UCSC genome browser and AceView. LD blocks comprise blocks of SNPs where all SNPs are in LD with *R*^2^ > 0.7.

| LD block | SNP | score* | CHR | BP | Band | VIP | Genes |
| --- | --- | --- | --- | --- | --- | --- | --- |
|  | rs17328231 | 5.96 | 1 | 95791119 | 1p31.1-q31.1 | 0.54 |  |
|  | rs6439445 | 4.24 | 3 | 135119256 | 3q12.3-q25.31 | 0.72 |  |
|  | rs16856619 | 4.08 | 3 | 146493435 | 3q12.3-q25.31 | 0.58 |  |
|  | rs345015 | 5.70 | 3 | 146667792 | 3q12.3-q25.31 | 0.62 |  |
| 1 | rs634364 | 4.18 | 6 | 53575551 | 6p21.1-q15 | 0.57 | AK126334, BC050580, AK125128, GCLC |
| 1 | rs525248 | 4.18 | 6 | 53576038 | 6p21.1-q15 | 0.57 | AK126334, BC050580, AK125128 |
| 2 | rs2148885 | 3.82 | 10 | 21413099 | 10p14-q24 | 0.57 | NEBL |
| 2 | rs11012530 | 5.06 | 10 | 21444536 | 10p14-q24 | 0.58 | NEBL |
| 3 | rs7897675 | 5.06 | 10 | 38448572 | 10p14-q24 | 0.55 | ZNF37A |
| 3 | rs17588142 | 5.06 | 10 | 38467714 | 10p14-q24 | 0.55 |  |
| 3 | rs7080636 | 5.06 | 10 | 38659180 | 10p14-q24 | 0.50 |  |
| 3 | rs34350622 | 5.17 | 10 | 41848403 | 10p14-q24 | 0.51 |  |
|  | rs12255371 | 5.15 | 10 | 41970728 | 10p14-q24 | 0.51 |  |
|  | rs7088870 | 3.91 | 10 | 73723319 | 10p14-q24 | 0.59 |  |
| 4 | rs7094314 | 3.83 | 10 | 82321942 | 10p14-q24 | 0.55 | SH2D4B |
| 4 | rs7904557 | 3.77 | 10 | 82326243 | 10p14-q24 | 0.56 | SH2D4B |
|  | rs12768174 | 5.82 | 10 | 84889167 | 10p14-q24 | 0.74 |  |
|  | rs10887866 | 4.48 | 10 | 90661730 | 10p14-q24 | 0.56 | STAMBPL1, KIAA1373, STAMBPL1andFAS |
|  | rs4646957 | 4.26 | 10 | 94219892 | 10p14-q24 | 0.54 | IDE |
|  | rs1235382 | 4.13 | 19 | 49711347 | 19p13.3-qter | 0.89 | CEACAM20 |

\* SNP-specific score indicating relative contribution to the RV statistic, as defined in text.

## 4 Discussion

In this report, we have taken a targeted approach to genetic association mapping of Alzheimer’s disease by focusing on SNPs in Alzheimer’s disease linkage regions and on imaging endophenotypes for brain regions affected by Alzheimer’s disease. We discovered association between SNPs in the linkage regions and the imaging endophenotypes, refined the set of SNPs by selecting those with high variable inclusion probabilities, and validated the refined set in an independent dataset. Here, we discuss our observations about the benefits and pitfalls of applying data-integration methods such as sparse canonical correlation analysis and the RV test in a high-dimensional data setting with low signal. We also discuss potential links between Alzheimer’s disease and genes in the priority set that were ranked highly in the validation data.

Initially, SCCA was used to find a subset of the SNPs in the linkage regions associated with the endophenotypes, but very little variable selection was achieved. SCCA uses a prediction criterion to identify the optimal soft-thresholding parameters for the sparse canonical variables, but using prediction error to select the penalty term includes irrelevant variables in the active set (Leng et al., 2006). In addition, the prediction-optimal value of the penalty term does not coincide with model selection consistency (Meinshausen and Buhlmann, 2006). Instead, we fixed the soft-thresholding parameter for the SNPs to achieve variable selection based on the rationale that no more than 7,500 SNPs (approximately 10%) are expected to be associated with the phenotypes. We applied bootstrapped-enhanced SCCA, a procedure analogous to the bootstrapped-enhanced elastic net proposed by Bunea et al. (2011) for imaging applications. To obtain a reduced set of SNPs to carry forward for validation, we thresholded the variable inclusion probabilities at 50%, as suggested by these authors, and at 90%. Bootstrapping to aid variable selection has been shown to be consistent in high-dimensional settings under some assumptions (Meinshausen and Buhlmann, 2010), and can improve recovery of the true model in regularized regression (Bach, 2008).

In a low-signal context, we do not necessarily expect to replicate association of the 22 “top-hit” SNPs in the validation data. For a fixed sample size, as the number of unassociated SNPs increases, the probability of a truly associated SNP being within the top-ranked SNPs decreases (Zaykin, 2005). By contrast, the more liberal threshold of *VIP* ≥ 50% resulted in a larger, “priority” set of 1,694 SNPs which could be validated and was substantially refined from the initial list of 75,845.

The permutation-based RV test of association proved to be a powerful tool in different phases of the analysis. This nonparametric test was computationally tractable and allowed us to uncover and validate linear association between the two multivariate datasets, one of them very high-dimensional, in an analysis setting with a low signal. Despite the evidence for association, the observed RV coefficient at each of the discovery, refinement and validation stages of the analysis was not large (< 0.1), consistent with SNPs having small association effects. The presence of SNPs with small effects is anticipated, as previous studies have found no large genetic effects apart from *APOE* (Ridge et al., 2013), for which we have already adjusted.

The ADNI studies use a case-control design, in which subjects are sampled conditional on meeting diagnostic criteria for either being cognitively normal, having late MCI, or having AD. Case-control designs do not result in a random sample from the population and they cannot be used to make inference about the population association between SNP genotypes and neuroimaging biomarkers without accounting for the biased sampling. To account for the biased sampling, we have applied inverse probability weighting in our analyses.

Investigation of the genes associated with the highest scoring SNPs in the validation data, reported in Table 3, identified genes previously implicated in AD. On chromosome 6, Glutamate-Cysteine Ligase Catalytic Subunit or *GCLC*, a gene annotation of the SNP rs634364, codes the first, rate-limiting enzyme of glutathione synthesis. Glutathione is an important antioxidant which plays an integrated role in the regulation of cell life, cell proliferation, and cell death (Pompella et al., 2003). The brain glutathione system is hypothesized to play a role in the breakdown of proteins in the brain, such as Aβ peptides (Lasierra-Cirujeda et al., 2013), and abundance of glutathione decreases with age and in some age-related disease (Liu et al., 2004). On chromosome 10, the complex locus *STAMPBL1andFAS* is an annotation of rs10887866 and codes a protein which plays a central role in programmed cell death (Choi and Benveniste, 2004). Through modulation of programmed cell death and neuronal atrophy, FAS may play a role in AD (Erten-Lyons et al., 2010). Also on chromosome 10, the gene insulin degrading enzyme (*IDE*) contains rs4646957 and codes the enzyme of the same name. *IDE* has previously been implicated in the progression Alzheimer’s disease as it degrades the Aβ peptides which are the main components in the amyloid plaques on the brains of subjects with Alzheimer’s disease (Edland et al., 2003). Edland et. al. found that three *IDE* variants were associated with risk of AD in subjects without copies of the ε4 *APOE* risk allele, the allele which constitutes the largest genetic risk of AD.

Gene expression from the UCSC RNA-Seq GTEx track was also explored to determine if any of the genes reported were highly expressed in the brain. On chromosome 10, Zinc Finger Protein 37A (*ZNF37A*), the gene containing rs7897675, is most highly expressed in the cerebellum and cerebellar hemisphere of the brain, regions related to motor function. Nebulette (*NEBL*), the gene annotation of rs2148885 and rs11012530, is most highly expressed in the heart, but has next highest gene expression in the brain. In addition, association fine-mapping under a linkage peak identified *NEBL* as a candidate gene for vitamin D levels in the blood (Aslibekyan et al., 2016). Low vitamin D blood levels are associated with accelerated decline in cognitive function in older adults (Miller et al., 2015).

Ten of the top 20 SNPs in Table 3 did not have associated gene annotations in the UCSC genome browser or AceView. For these SNPs, flanking genes were queried with ALFRED (Rajeevan et al., 2011) and the UCSC genome browser, since SNPs may “tag” causal variants in nearby genes. Genes were considered to be flanking if they were within 1 Mb of the SNPs in the priority set, though many of the flanking genes reported are much closer to the priority SNPs. On chromosome 3, rs643944 is approximately 22 kb proximal to the flanking gene *RAB6B*. *RAB6B* is the brain-specific isoform of *RAB6* (Wanschers et al., 2007), a family of proteins which impair the processing of the amyloid precursor protein involved in the development of AD (*Thyrock et al., 2013*). On chromosome 10, *DDIT4* is approximately 17.5 kb proximal to rs7088870. *DDIT4* produces the REDD1 protein, which enhances stress-dependent neuronal cell death and is involved in dysregulation of the mammalian target of rapamycin (mTOR) pathway (Maiese, 2014). Dysregulation of mTOR is a hallmark of a wide variety of brain disorders (Polman et al., 2012), and inhibition of mTOR is associated with Aβ-peptide-related synaptic dysfunction in AD (Ma et al., 2010). Another flanking gene to rs7088870 is *DNAJB12*, which is approximately 39.2 kb proximal to rs7088870, and is involved in protein folding. The process of plaque buildup in AD involves the accumulation of misfolded Aβ proteins, and *DNAJB12* is highly expressed throughout the brain (Tebbenkamp and Borchelt, 2010). Finally, in addition to being the gene annotation of rs7897674, *ZNF37A* is also 15.4 kb proximal to the SNP rs17588142 on chromosome 10.

In summary, this analysis illustrates the application of novel methods for integration of high-dimensional data with low signal. To focus on regions with increased prior probability of containing deleterious variants, the analysis was restricted to SNPs within linkage regions for AD. In practice, the same methodology could be applied to all available data if no linkage regions have previously been identified. However, we recommend restricting analyses to linkage regions whenever possible to leverage information from prior work. The objective was to obtain a refined list of SNPs to propose for further investigation. Naive application of SCCA did not lead to any refinement, potentially due to the data containing many small effects. Instead, we were able to obtain refinement through bootstrapped-enhanced SCCA. Throughout, the analysis benefited from the RV test to assess the evidence of linear association between two multivariate datasets: the high-dimensional genomic data, and the multidimensional neuroimaging data. RV tests of SNPs selected based on variable importance probabilities identified a priority set of 1,694 SNPs in the ADNI-1 data that was associated with the rates of changes in the brain regions of interest in the ADNI-2 validation set. Our final results are encouraging, in that genes corresponding to SNPs with the highest contributions to the RV coefficient in the validation data have previously been implicated or hypothesized to be implicated in AD, including *GCLC*, *IDE*, and *STAMBP1andFAS*. We hypothesize that the effect sizes of the 1,694 SNPs in the priority set are likely small, but further investigation within this set may advance understanding of the missing heritability in late-onset Alzheimer’s disease.

## 5 Acknowledgements

Data collection and sharing was funded by the Alzheimer’s Disease Neuroimaging Initiative (ADNI) (National Institutes of Health Grant U01 AG024904) and DOD ADNI (Department of Defense award number W81XWH-12-2-0012). ADNI is funded by the National Institute on Aging, the National Institute of Biomedical Imaging and Bioengineering, and through generous contributions from AbbVie, Alzheimer’s Association; Alzheimer’s Drug Discovery Foundation; Araclon Biotech; BioClinica, Inc.; Biogen; Bristol-Myers Squibb Company; CereSpir, Inc.; Eisai Inc.; Elan Pharmaceuticals, Inc.; Eli Lilly and Company; EuroImmun; F. Hoffmann-La Roche Ltd and its affiliated company Genentech, Inc.; Fujirebio; GE Healthcare; IXICO Ltd.; Janssen Alzheimer Immunotherapy Research & Development, LLC.; Johnson & Johnson Pharmaceutical Research & Development LLC.; Lumosity; Lundbeck; Merck & Co., Inc.; Meso Scale Diagnostics, LLC.; NeuroRx Research; Neurotrack Technologies; Novartis Pharmaceuticals Corporation; Pfizer Inc.; Piramal Imaging; Servier; Takeda Pharmaceutical Company; and Transition Therapeutics. The Canadian Institutes of Health Research is providing funds to support ADNI clinical sites in Canada. Private sector contributions are facilitated by the Foundation for the National Institutes of Health (<u>www.fnih.org</u>). The grantee organization is the Northern California Institute for Research and Education, and the study is coordinated by the Alzheimer’s Disease Cooperative Study at the University of California, San Diego. ADNI data are disseminated by the Laboratory for Neuro Imaging at the University of Southern California.

This work is based on Elena Szefer’s MSc thesis supervised by J Graham and was supported in part by the Natural Sciences and Engineering Research Council of Canada. The authors would like to thank Ellen Wijsman for helpful discussions about *APOE* and population stratification and Wayne Wang for assistance with the genomic quality control of the ADNI-2 validation data. The authors are grateful to the anonymous reviewers for constructive comments which greatly improved the manuscript.

---

**Table 1A.**
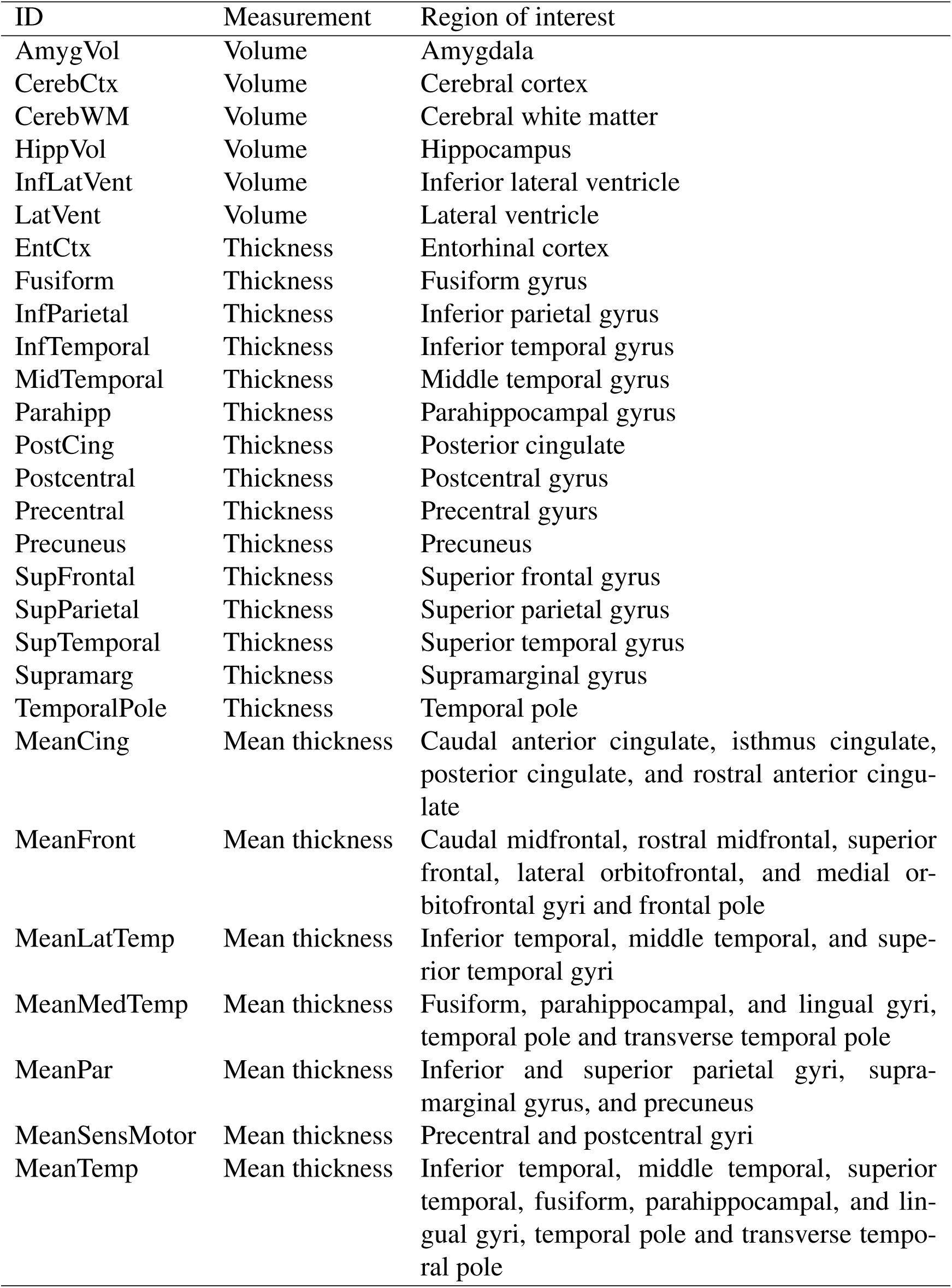
Imaging phenotypes defined as volumetric or cortical thickness measures of 28 × 2 = 56 regions of interest (ROIs) from automated Freesurfer parcellations. Each of the phenotypes in the table corresponds to two phenotypes in the data: one for the left hemisphere and the other for the right hemisphere.

